# Understanding the interaction of Hsp70 with NK cells

**DOI:** 10.1101/2023.07.27.550847

**Authors:** Ruchir Sahni, Gabriele Multhoff

## Abstract

Heat Shock Protein 70 (Hsp70) is a highly conserved and ubiquitous molecular chaperone that plays a central role in cellular protein machinery and stress response. Membrane-bound HSP70 has emerged as an important cancer biomarker and acts as a danger signal and elicits immune response. Hsp70 membrane expression is correlated to increased sensitivity to lysis carried out by NK cells. This study uses computational approaches to decode the interaction of Hsp70 with NK cells and determines the binding site for Hsp70 on the surface of NK cells. Our findings identified CD69 and NKP46 as the most probable binding sites for Hsp70. Additionally, we confirmed the strong binding affinity between Hsp70 and the CD94-NKG2A complex.

## 1 Introduction

Heat shock proteins (HSPs) are proteins that are found universally and are constitutively expressed in human cells. The production of these molecules within cells is triggered by different types of “stress signals,” such as elevated temperature exposure, ultraviolet (UV) light exposure, reactive oxygen species, metals with high density, cytostatic medications, as well as viral or bacterial invasions. HSPs are molecular chaperones which are involved in protein folding/defolding, protein transport and protection mechanisms from protein aggregration (Hartl & Hayer-Hartl 2002). Within tumor cells, Hsp70 plays a critical role in promoting tumor cell survival by disrupting apoptosis pathways (Jäättelä 1999; Jäättelä et al. 1998).

In addition to its intracellular chaperoning functions, HSPs have been observed to be expressed on the outer cell surface of highly aggressive primary and metastatic tumor cells (Ferrarini et al. 1992). Extracellularly localised HSPs serve dual roles: they can act as carrier molecules, facilitating the transport of immunogenic peptides to antigen presenting cells (APCs) (Tamura et al. 1997; Schild et al. 1999; Udono & Srivastava 1993) or they can directly stimulate the innate immune system by secreting pro-inflammatory cytokines (Asea et al. 2000a) (Asea et al. 2000b). It was demonstrated that Hsp70 is the recognition site for cytosolic attack by NK cells (Multhoff et al. 1997). Antibody blocking studies revealed that Hsp70 is needed for the recognition by transiently plastic adherent NK cells (Multhoff et al. 1995, 1997; Botzler et al. 1998). Furthermore, specific monoclonal antibodies (mAbs) targeting Hsp70 have been shown to block the cytolytic activity of NK cells, indicating the significance of Hsp70 in regulating NK cell-mediated cytotoxicity (Multhoff et al. 1995).

The interaction between Hsp70 and NK cells is not well understood. CD 94 is predicted to play an important role in the interaction (Gross et al. 2003a,b) but it needs to be studied further. Interestingly, a 14-mer peptide TKDNNLLGRFELSG (TKD, aa 450-463) part of Hsp70 induces the cytolytic and proliferative activity of NK cells at concentrations equivalent to full-length Hsp70 protein (Multhoff et al. 2001). This study aims to find the binding site of Hsp70 on NK cell surface. We perform molecular docking of Hsp70 with multiple NK cell surface receptors using protein-protein docking algorithms. We also performed molecular docking by tagging the TKD peptide as the active site of the interaction. Then, binding energies and interactions of complexes were calculated. We additionally performed molecular docking on CD69-NKG2A complex (PDB ID : 3BDW) as previous studies predict CD94 to play an important role in the interaction (Gross et al. 2003a,b; Brooks et al. 1997).

## 2 Materials and Methods

### Protein Preparation

The following table (Table 1) lists the receptors used in the study. These receptors are the most common receptors found on the surface of NK cells.

**Table 1:**
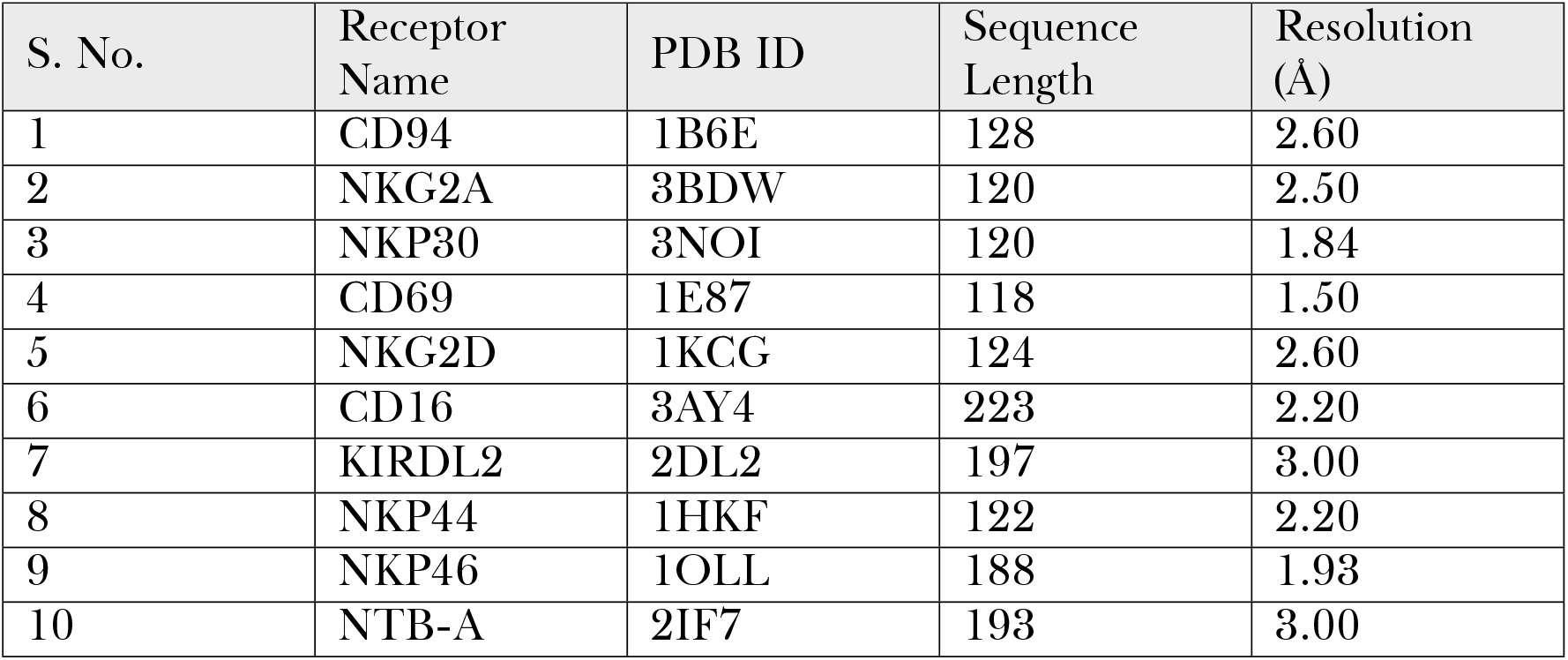
List of NK receptors used in the study.

| S. No. | Receptor Name | PDB ID | Sequence Length | Resolution (Å) |
| --- | --- | --- | --- | --- |
| 1 | CD94 | 1B6E | 128 | 2.60 |
| 2 | NKG2A | 3BDW | 120 | 2.50 |
| 3 | NKP30 | 3NOI | 120 | 1.84 |
| 4 | CD69 | 1E87 | 118 | 1.50 |
| 5 | NKG2D | 1KCG | 124 | 2.60 |
| 6 | CD16 | 3AY4 | 223 | 2.20 |
| 7 | KIRDL2 | 2DL2 | 197 | 3.00 |
| 8 | NKP44 | 1HKF | 122 | 2.20 |
| 9 | NKP46 | 1OLL | 188 | 1.93 |
| 10 | NTB-A | 2IF7 | 193 | 3.00 |

The PDB files of the receptor were downloaded using RCSB website (https://www.rcsb.org/). For submission of files into the docking softwares, the structures cannot have more than one chain, cannot have overlapping residue numbers, and cannot have alternate locations for any atoms. PDBTOOLS (https://wenmr.science.uu.nl/pdbtools/) was used to produce these files (Jiménez-García et al. 2021). Next, heteroatoms such as solvent molecules and ligands were removed during protein preparation in PDBTOOLS. Further, the missing hydrogens in protein structures were added using ChimeraX (Pettersen et al. 2021).

### Molecular Docking

Each receptor is docked to Hsp70 (PDB ID : 4PO2) by employing two softwares : Cluspro 2.0 (https://cluspro.org/) and Haddock 2.4 (https://wenmr.science.uu.nl/haddock2.4/). First, the receptor and Hsp70 pdb file is uploaded to Cluspro 2.0 to get the most favourable docking structure. ClusPro 2.0 is an online platform designed for the direct docking of two interacting proteins. This web server executes three computational steps, outlined as follows: (1) rigid body docking, involving the sampling of billions of conformations; (2) root-mean-square deviation (RMSD) based clustering of the 1000 lowest energy structures to identify the largest clusters, representing the most probable models of the complex; and (3) refinement of selected structures through energy minimization. ClusPro provides the top 30 complexes for each job following CHARMM energy minimization and considering both clustering probability and energy-based parameters (balanced, electrostatic-favored, hydrophobic-favored, and VDW+electrostatic-favored) (Kozakov et al. 2017). In this study, the best complexes were selected based on the balanced energy parameter and the top cluster was chosen.

Then, the receptor and Hsp70 pdb files is uploaded to Haddock 2.4 and TKD peptide is marked as active residue on Hsp70 while the entire receptor is marked as the active residue on the receptor. HADDOCK utilizes biochemical and/or biophysical interaction data, such as chemical shift perturbation data from NMR titration experiments or mutagenesis data. The data on the interacting residues is incorporated as ambiguous interaction restraints (AIRs) to guide the docking process. Following the calculation, the generated structures are ranked based on their intermolecular energy, which comprises the sum of electrostatic, van der Waals, and AIR energy terms (van Zundert et al. 2016). In this study, the structure with the lowest Haddock Score is chosen.

### Calculation of Energetics

These docking complexes were then analysed by calculating energetics of the interaction. To calculate the binding affinity of the complex, PRODIGY web tool (https://wenmr.science.uu.nl/prodigy/) was used. The results encompass the predicted binding free energy value (ΔG) expressed in kcal mol^−1^, as well as the projected dissociation constant (Kd) in M, determined using the equation ΔG = RT ln(Kd), where R signifies the ideal gas constant (kcal K-1*mol*^−1^), and T represents the temperature (K) (Xue et al. 2016).

PPCheck (http://caps.ncbs.res.in/ppcheck/) was used to measure the energy calculations of the interaction. PPCheck is a web server designed to assess the strength of interactions between any two provided proteins or chains and its results include H-bond energy, Electrostatic energy, van der Waals energy and Total Stabilizing Energy (Sukhwal & Sowdhamini 2013).

## 3 Results

Here, we dock Hsp70 with each of the receptor in two different ways : 1) Docking and selecting results with best score (using Cluspro 2.0) 2) Docking while keeping TKD as the active residue (using Haddock 2.4) (Figure 1).

**Figure 1.**
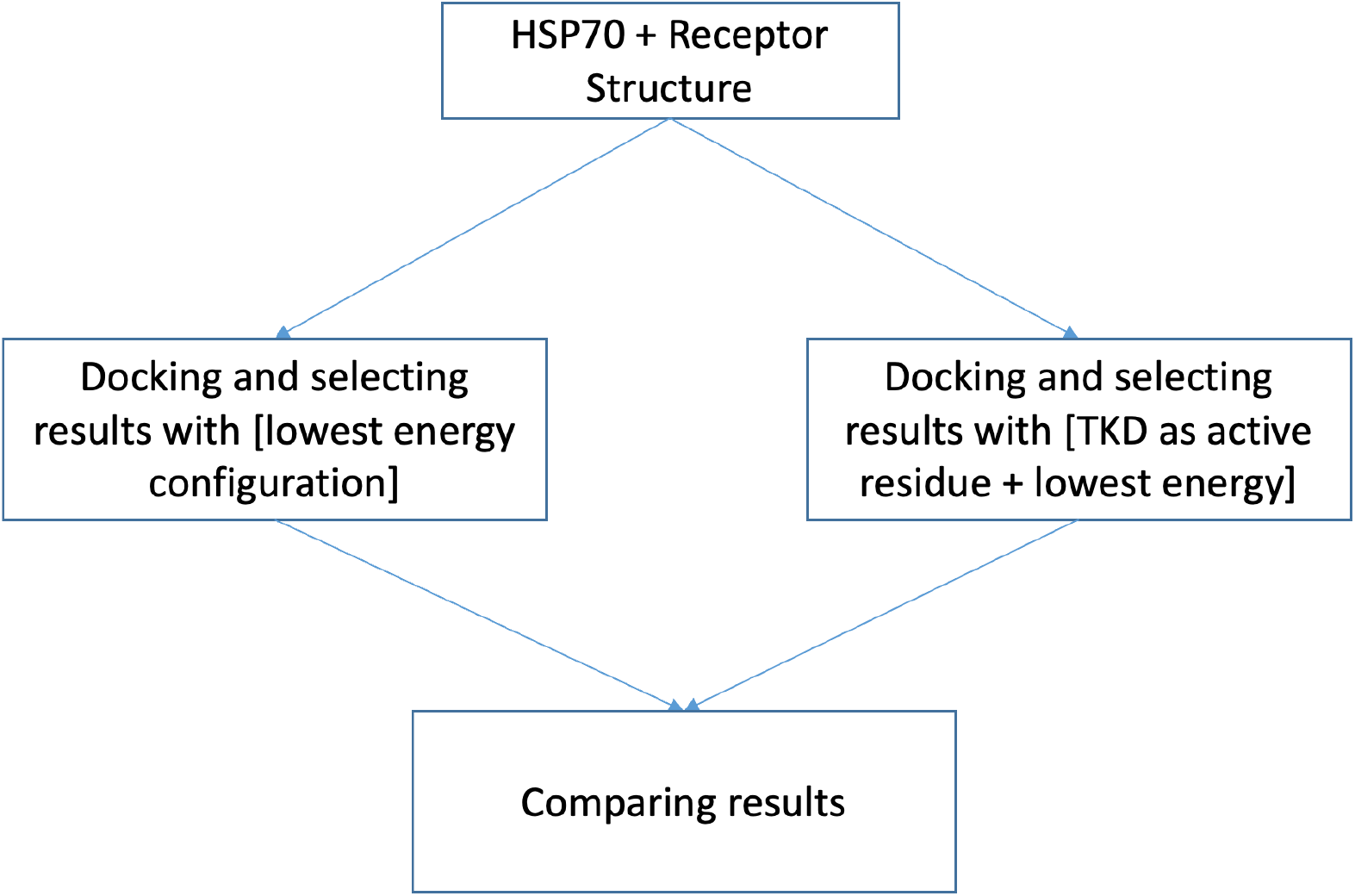
: Workflow of the docking performed in this study

The results of docking from Cluspro 2.0 software are shown in Figure 2. The figures were labelled as : Hsp70 -Purple, Receptor -Green and TKD peptide -Red. The figures are labelled and prepared using ChimeraX.

**Figure 2.**
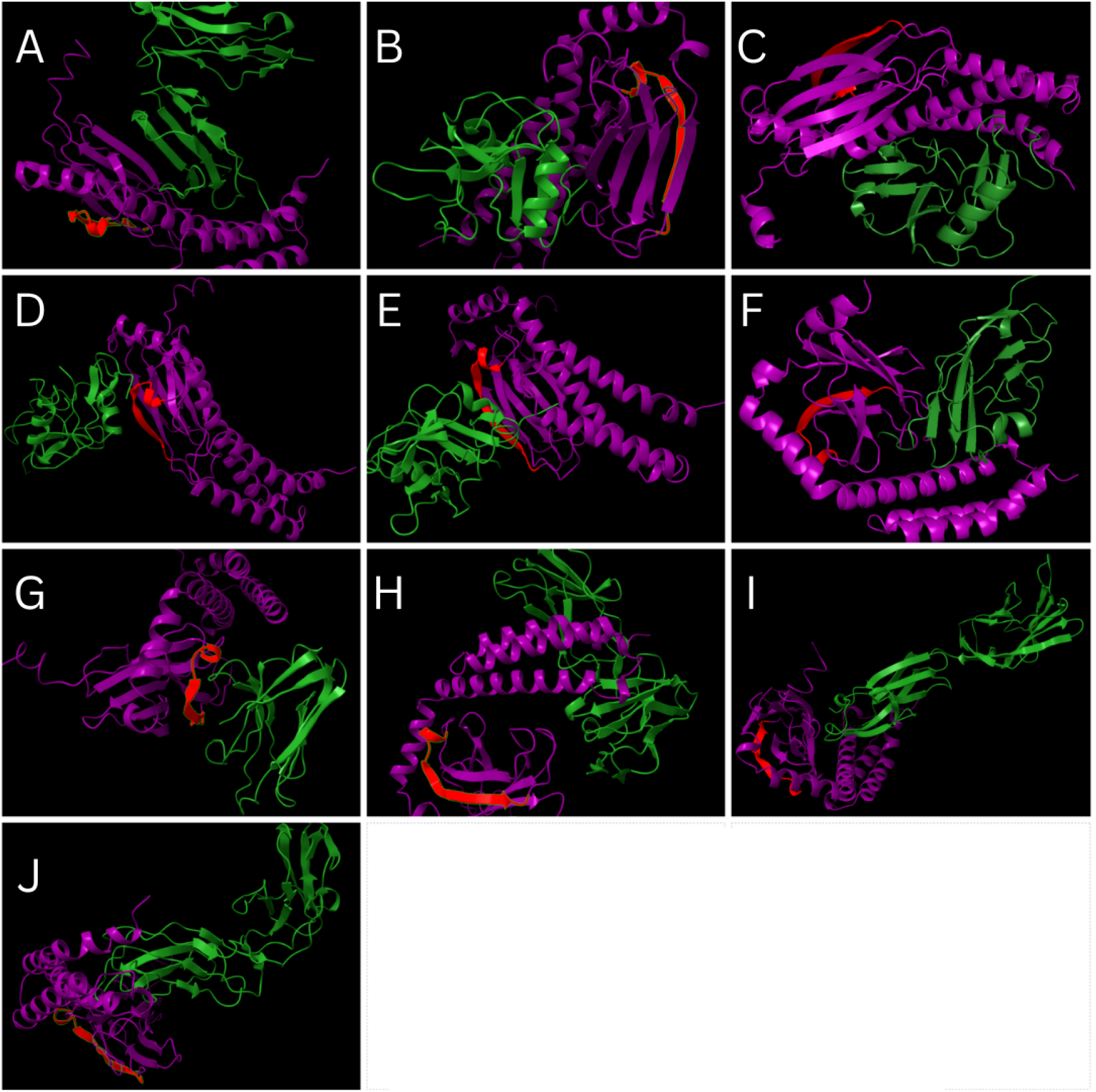
: Results of docking from Cluspro 2.0. A -CD16, B -CD69, C -CD94, D -NKG2A, E -NKG2D, F -NKP30, G -NKP44, H -NKP46, I -NTB-A, J -KIRDL2

The results of docking from Haddock 2.4 software are shown in Figure 3. The figures were labelled as : Hsp70 -Purple, Receptor -Green and TKD peptide -Red. The figures are labelled and prepared using ChimeraX.

**Figure 3.**
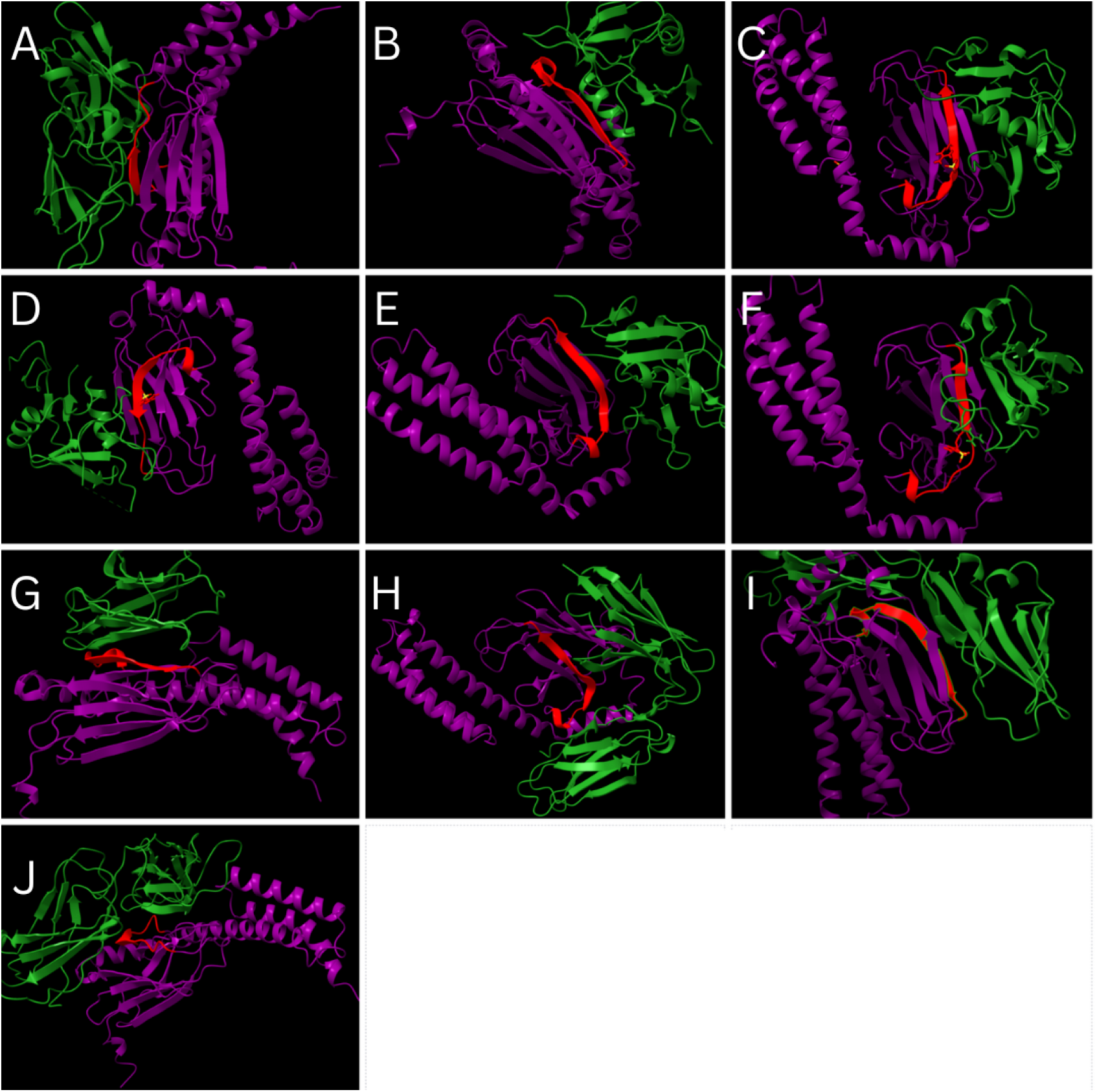
: Results of docking from Haddock 2.4 (TKD tagged as active residue). A -CD16, B -CD69, C -CD94, D -NKG2A, E -NKG2D, F -NKP30, G -NKP44, H -NKP46, I -NTB-A, J -KIRDL2

**Figure 4.**
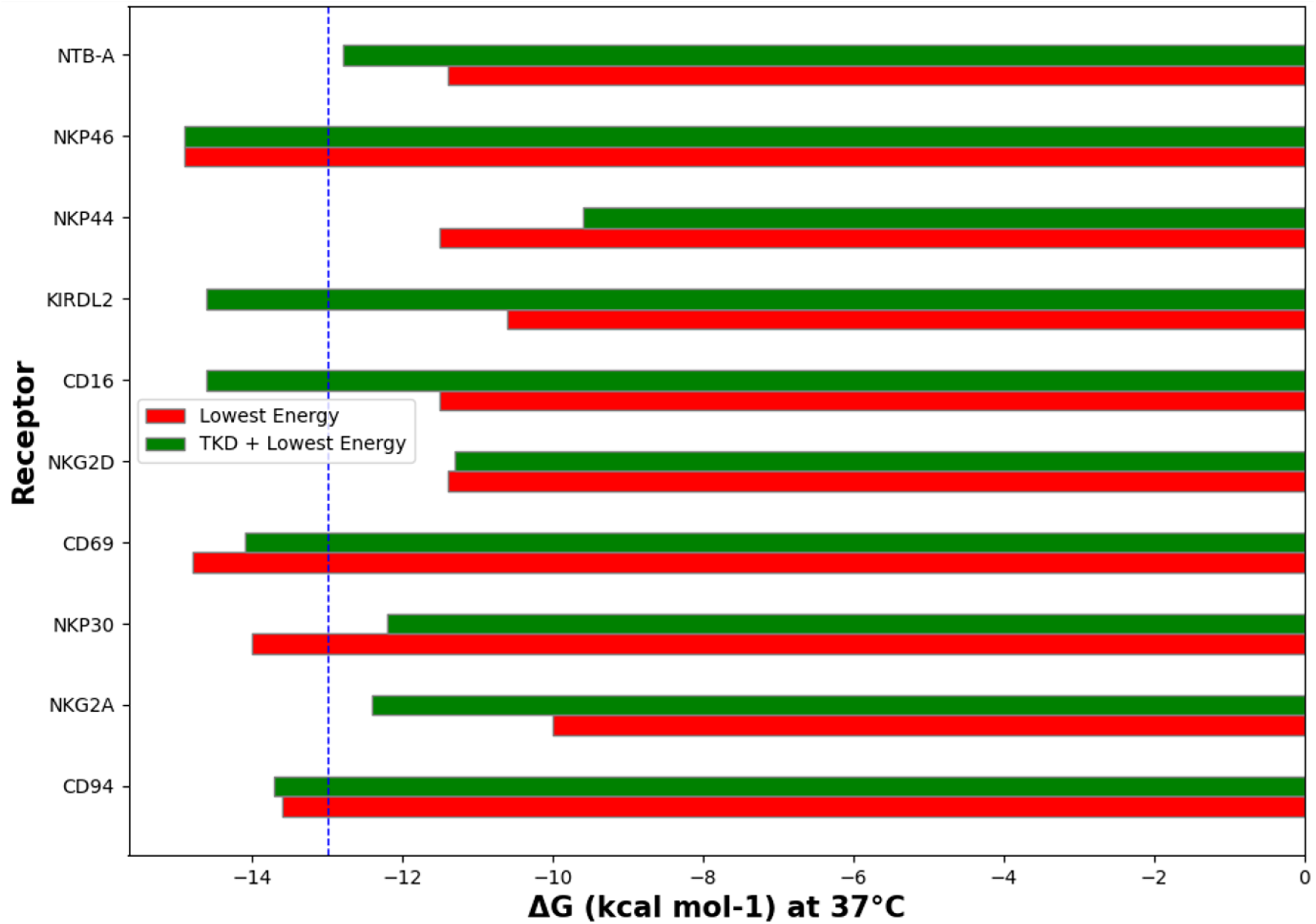
: Binding energies of different receptor-Hsp70 complex

### Among individual receptors, CD69 and NKP46 are the most probable binding site for HSP70

Binding energies of each complex calculated by Prodigy web-server is displayed on Figure 5. Among the 10 complexes, the binding energies vary from -9.6 to -14.9 kcal mol-1. It is noted that CD69 and NKP46 have the most binding energy among the individual receptors and can be considered the most probable binding sites. Surprisingly, KIRDL2 and CD16 show a significant increase in binding energy in complexes from Haddock 2.4 (where TKD is tagged as active residue) compared to the results from Cluspro 2.0.

**Figure 5.**
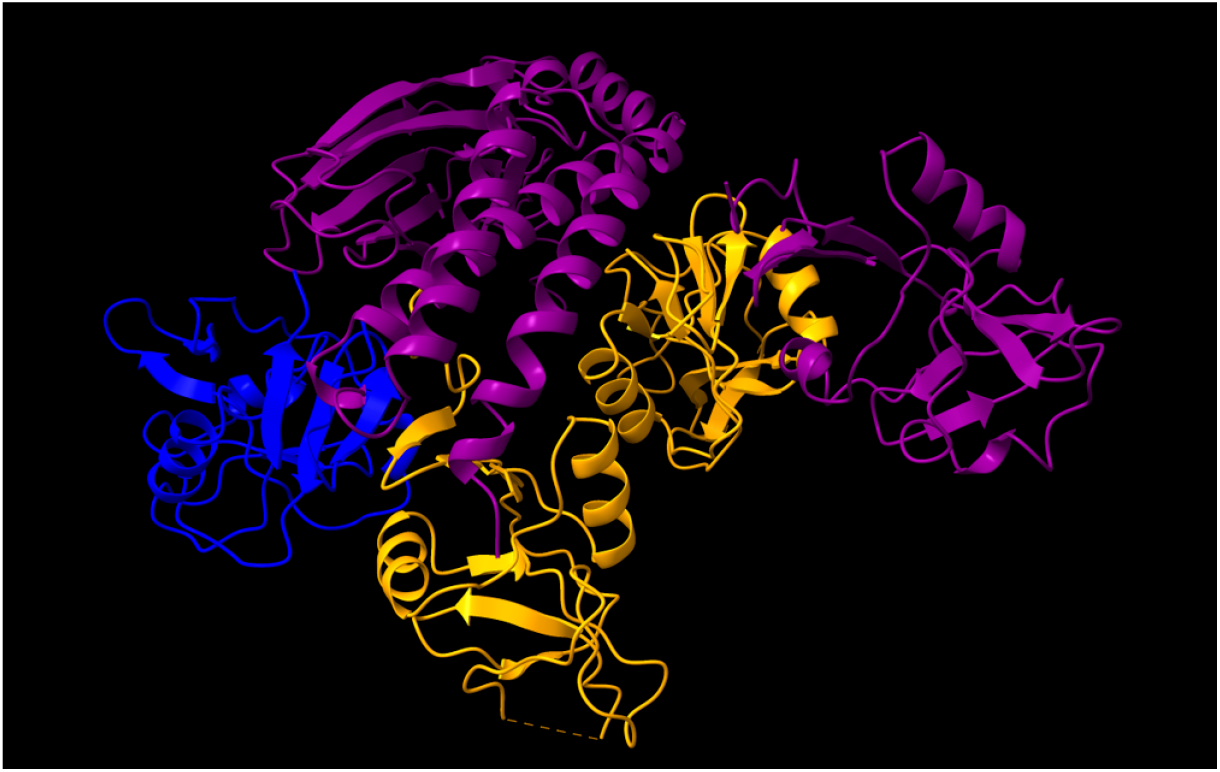
: Results of docking of CD94-NKG2A complex with Hsp70 using Cluspro 2.0. Purple -Hsp70, Blue -CD94, Yellow -NKG2A

To further understand the interaction between the complexes, PPCheck webserver was used to calculate the energetics of the interaction. The calculations for CD69, NKP46, KIRDL2 and CD16 are shown in Table 2.

**Table 2:**
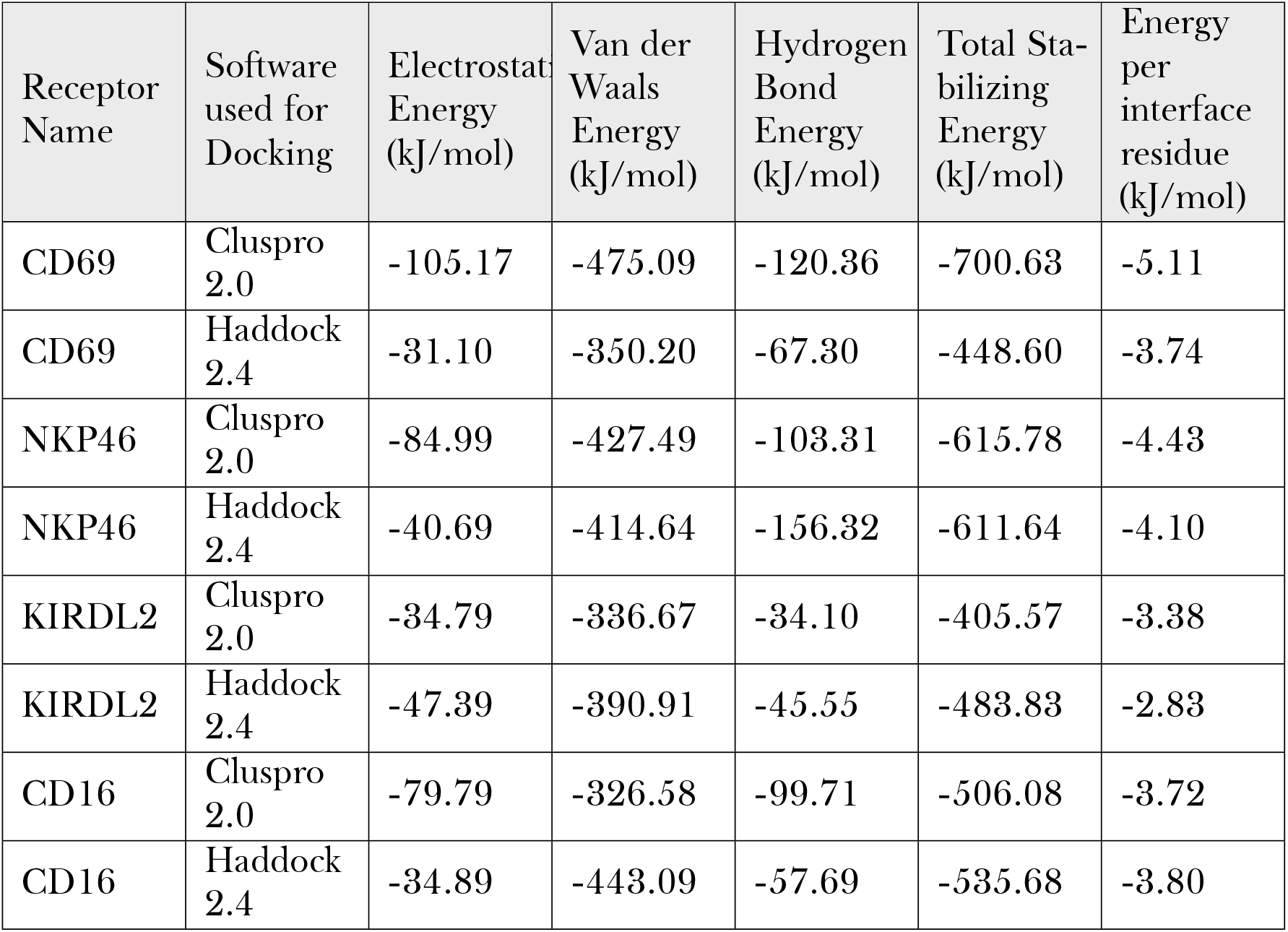
Energetics of the complexes.

| Receptor Name | Software used for Docking | Electrostatic Energy (kJ/mol) | Van der Waals Energy (kJ/mol) | Hydrogen Bond Energy (kJ/mol) | Total Stabilizing Energy (kJ/mol) | Energy per interface residue (kJ/mol) |
| --- | --- | --- | --- | --- | --- | --- |
| CD69 | Cluspro 2.0 | -105.17 | -475.09 | -120.36 | -700.63 | -5.11 |
| CD69 | Haddock 2.4 | -31.10 | -350.20 | -67.30 | -448.60 | -3.74 |
| NKP46 | Cluspro 2.0 | -84.99 | -427.49 | -103.31 | -615.78 | -4.43 |
| NKP46 | Haddock 2.4 | -40.69 | -414.64 | -156.32 | -611.64 | -4.10 |
| KIRDL2 | Cluspro 2.0 | -34.79 | -336.67 | -34.10 | -405.57 | -3.38 |
| KIRDL2 | Haddock 2.4 | -47.39 | -390.91 | -45.55 | -483.83 | -2.83 |
| CD16 | Cluspro 2.0 | -79.79 | -326.58 | -99.71 | -506.08 | -3.72 |
| CD16 | Haddock 2.4 | -34.89 | -443.09 | -57.69 | -535.68 | -3.80 |

**Table 3:**
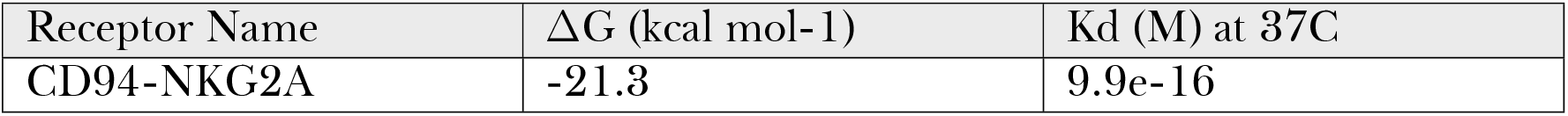
Binding Affinity of Hsp70 with the complex CD94-NKG2A.

| Receptor Name | $\Delta G$ (kcal mol<sup>-1</sup>) | Kd (M) at 37C |
| --- | --- | --- |
| CD94-NKG2A | -21.3 | 9.9e-16 |

The energetics calculations show that the difference of binding affinity in CD16 and KIRDL2 between the two conditions is due to the difference in the total stabilizing energy. CD69 and NKP46 have the lowest total stabilizing energy and hence results in highest binding affinity. These complexes also have the lowest energy per interface residue, making the complexes very stable.

### Hsp70 binds strongly to the CD94-NKG2A complex

Previous literature (Gross et al. 2003a,b; Brooks et al. 1997) has shown that CD94-NKG2 complex plays an important role in the activation of NK cells by Hsp70. It is noted that the binding affinity of Hsp70 with CD94-NKG2A complex is higher that the binding affinity of Hsp70 with any of the individual receptors.

## 4 Discussion

In this study, our objective was to understand the interaction between Heat Shock Protein 70 (Hsp70) and Natural Killer (NK) cells using computational approaches.Hsp70 is a highly conserved molecular chaperone that plays a crucial role in cellular protein machinery and stress response. Previous investigations have indicated that Hsp70, when localized on the cell surface, acts as a cancer biomarker and serves as a danger signal, triggering an immune response. Additionally, its membrane expression has been correlated with increased sensitivity to NK cell-mediated lysis.

Hsp70 is not solely involved in intracellular chaperoning functions; it has also been observed to be expressed on the outer cell surface of aggressive primary and metastatic tumor cells. Extracellularly localized Hsp70 exhibits two distinct roles: 1) it acts as a carrier molecule, facilitating the transport of immunogenic peptides to antigen-presenting cells (APCs), and 2) it can directly stimulate the innate immune system by secreting pro-inflammatory cytokines. Moreover, Hsp70 has been identified as a recognition site for cytosolic attack by NK cells, further highlighting its significance in regulating NK cell-mediated cytotoxicity.

In this study, we performed molecular docking simulations of Hsp70 with multiple NK cell surface receptors to identify potential binding sites. For the docking process, we utilized two software tools, namely ClusPro 2.0 and Haddock 2.4 for two distinct protocols : Haddock 2.4 tagged TKD peptide as an active residue while ClusPro 2.0 selected results without such specification. Among the receptors under study, CD69 and NKP46 emerged as the most probable binding sites for Hsp70.

To gain insights into the stability of the interactions, we calculated the binding energies of the complexes using the Prodigy web tool and PPCheck web server. The CD69 and NKP46 complexes exhibited the lowest total stabilizing energy, indicating high stability and a strong binding affinity with Hsp70. Interestingly, the difference in binding affinity observed between the CD16 and KIRDL2 complexes in the two docking conditions (ClusPro 2.0 and Haddock 2.4) was attributed to variations in the total stabilizing energy.

Additionally, we investigated the interaction between Hsp70 and the CD94-NKG2A complex. Prior studies have highlighted the role of the CD94-NKG2 complex in NK cell activation by Hsp70. In our study, we found that the binding affinity of Hsp70 with the CD94-NKG2A complex was higher than its binding affinity with any of the individual receptors, further supporting the significance of this complex in the interaction.

In conclusion, our computational approaches provided valuable insights into the interaction between Hsp70 and NK cells. We identified potential binding sites on NK cell surface receptors and found that CD69 and NKP46 are the most probable binding sites for Hsp70. Moreover, we confirmed the strong binding affinity between Hsp70 and the CD94-NKG2A complex. These findings contribute to a better understanding of the molecular mechanisms governing Hsp70-NK cell interactions and lay the groundwork for future studies exploring therapeutic interventions targeting Hsp70 in cancer immunotherapy.

## 5 Acknowledgements

We are grateful to Technische Universität München for providing access to necessary resources and facilities during the research.

